# Extinction Training Suppresses Alcohol Relapse by Inhibiting Acquisition-Recruited Striatal Ensembles and Engaging Striosomal Neurons

**DOI:** 10.64898/2026.04.30.722024

**Authors:** Xueyi Xie, Xuehua Wang, Jun Wang

## Abstract

Relapse to alcohol use is frequently triggered by re-exposure to alcohol-associated cues, and extinction-based interventions reduce relapse vulnerability. However, the cellular mechanisms through which extinction suppresses alcohol seeking remain unclear. Using a mouse model of operant alcohol self-administration, we examined how extinction training alters activity of defined striatal direct-pathway medium spiny neuron (dMSN) populations in the dorsomedial striatum (DMS). Extinction significantly reduced cue-induced reinstatement and decreased reactivation of acquisition-recruited dMSN ensembles during relapse. Chemogenetic activation of acquisition-recruited dMSN ensembles during extinction impaired extinction learning and enhanced subsequent reinstatement, even in the absence of manipulation at test, indicating that suppression of these ensembles is required for effective extinction. In contrast, selective activation of striosomal dMSNs during extinction accelerated extinction learning and further reduced reinstatement without affecting locomotor activity. These findings demonstrate that extinction suppresses alcohol seeking through coordinated modulation of distinct striatal dMSN populations, involving both reduced engagement of acquisition-related ensembles and recruitment of striosomal circuits. Together, these findings provide mechanistic insight into how extinction reshapes striatal circuitry to suppress relapse-related behavior and highlight defined striatal dMSN populations as potential substrates for enhancing extinction-based interventions.

## Introduction

Relapse remains one of the greatest challenges in the treatment of alcohol and substance use disorder^1–3^. Even after prolonged abstinence, exposure to drug-associated cues can trigger renewed drug seeking, reflecting the persistence of maladaptive drug-related memories^1,2,4^. In both clinical and preclinical settings, extinction-based interventions have been shown to reduce relapse vulnerability^5–10^. However, extinction does not erase the original memory; rather, relapse can re-emerge under stress, context shifts, or cue exposure^9,11–13^. Although extinction training reduces relapse probability, the neural mechanisms by which extinction suppresses alcohol seeking remain incompletely understood. Elucidating the cellular mechanisms underlying extinction is therefore critical for developing strategies to enhance extinction efficacy and more effectively prevent relapse.

At a conceptual level, extinction is thought to involve at least two complementary processes^11^. First, extinction may suppress the expression of previously acquired drug memories. Second, extinction represents a form of new learning that encodes updated action–outcome contingencies. How these processes are implemented at the level of defined neuronal ensembles remains a central unanswered question in addiction neuroscience.

The dorsomedial striatum (DMS) plays a critical role in goal-directed action and alcohol seeking^5,14–16^. Within the striatum, direct-pathway medium spiny neurons (dMSNs) promote action execution and reinforcement learning^17–22^. Recent evidence, including our prior work, indicates that subsets of DMS dMSNs recruited during acquisition of operant alcohol self-administration form ensembles that encode alcohol-associated memories^23–27^. Reactivation of these acquisition-recruited dMSN ensembles during cue exposure drives relapse-like behavior, suggesting that persistent striatal activity patterns contribute directly to reinstatement of alcohol seeking.

If acquisition-recruited dMSN ensembles encode alcohol memories that drive relapse, one possibility is that extinction suppresses relapse by reducing the reactivation of these ensembles. In this model, extinction training would dampen the influence of acquisition-encoded striatal representations, thereby limiting cue-induced alcohol seeking. However, direct evidence that extinction alters the re-engagement of acquisition-recruited striatal ensembles during relapse is lacking. Moreover, whether suppression of these ensembles is necessary for successful extinction learning remains unknown.

In addition to suppressing previously formed memory representations, extinction is widely considered an active learning process^9,12,28^. The striatum is not a homogeneous structure but is organized into matrix and striosome compartments that differ in connectivity, neurochemical identity, and functional roles^29–32^. Emerging studies suggest that in contrast to matrix dMSNs which drive positive reinforcement and behavioral execution, striosomal dMSNs may be preferentially involved in negative reinforcement and behavioral inhibition^29,32–37^. These observations raise the possibility that extinction engages distinct striatal subpopulations in the striosome to constrain alcohol seeking.

Here, we investigated how extinction training suppresses cue-induced reinstatement of alcohol seeking at the level of defined striatal dMSN populations. We first tested whether extinction reduces reactivation of acquisition-recruited dMSN ensembles during reinstatement. We then examined whether maintaining activity of these ensembles during extinction using DREADDs (Designer Receptors Exclusively Activated by Designer Drugs) interferes with extinction learning. Finally, we asked whether enhancing activity of striosomal dMSNs during extinction facilitates extinction and confers more persistent protection against relapse.

## Results

### Extinction training suppressed cue-induced reinstatement of alcohol-seeking behavior

Previous studies showed that extinction training reduces cue- or context-induced reinstatement of other drug-seeking behaviors^8,38–40^, we thus first asked whether operant alcohol extinction suppresses cue-induced reinstatement of alcohol seeking. Wildtype mice were trained in an operant alcohol self-administration paradigm for ∼8 weeks (Fig. 1A). The chamber contained retractable active and inactive levers flanking a central magazine, with a cue light positioned above the magazine. Under a fixed ratio 1 (FR1) schedule, each active lever press resulted in alcohol delivery paired with a cue light and tone for 20 sec. During this period, additional lever presses were recorded but had no programmed consequences. Training progressively advanced to FR2 and then FR3. Mice achieved stable responding during the final FR3 sessions (Fig. 1B, left).

**Figure 1.**
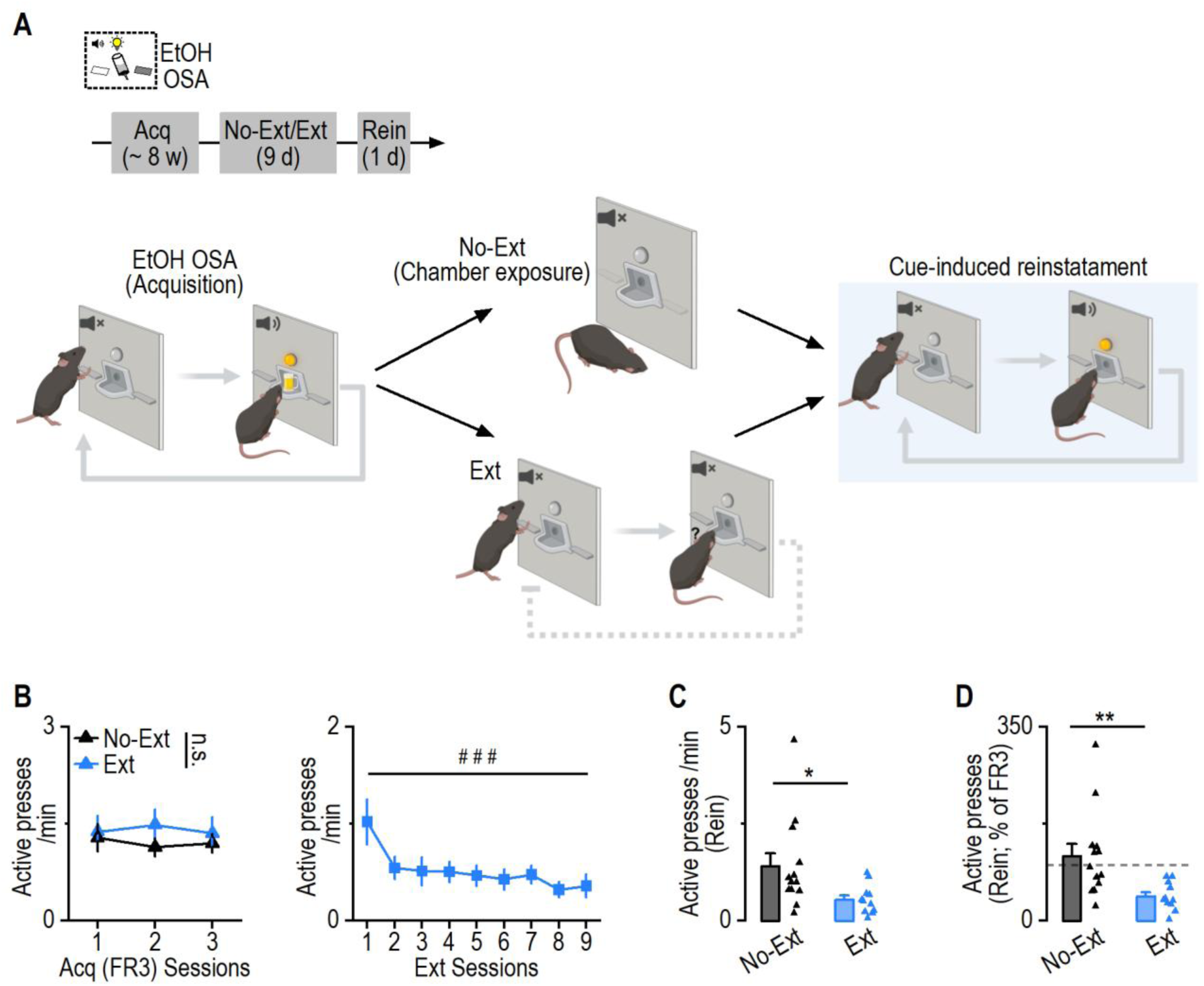
Extinction training suppressed cue-induced reinstatement of alcohol seeking. ***A*,** Schematic of the behavioral paradigm. Mice underwent acquisition of operant alcohol self-administration (Acq; ∼8 weeks), followed by either extinction training (Ext; 9 days) or no-extinction control (No-Ext). All groups were subsequently tested for cue-induced reinstatement. ***B*,** Left: Active lever pressing during the final acquisition sessions (FR3 schedule) was comparable between groups prior to extinction or no-extinction manipulation. Two-way RM ANOVA: F_(1,22)_ = 0.579, p = 0.455. Right: Active pressing gradually reduced during 9-d extinction training for the extinction group. Friedman’s test: between sessions: X^2^ = 28.13, ^###^p = 0.0005. ***C*,** Mice with a history of extinction exhibited reduced active lever pressing during reinstatement compared to no-extinction controls. Mann-Whiteney test: U = 29.5, *p = 0.013. ***D*,** Reinstatement responding expressed as a percentage of acquisition responding demonstrates that extinction training significantly suppressed alcohol relapse. Mann-Whiteney test: U = 22, **p = 0.0031. n = 13 mice (No-Ext) and 11 mice (Ext).

Following acquisition, mice were divided into two groups. One group underwent 9 days of lever extinction, during which active lever presses no longer produced alcohol or cue presentations, leading to a gradual reduction in responding (Fig. 1B, right). The control group was returned daily to the chamber for the same duration but with levers retracted, controlling for context exposure without actual lever-extinction learning.

One day after the final extinction session, both groups underwent a cue-induced reinstatement test. Active lever presses were significantly lower in the extinction group compared to controls (Fig. 1C). When reinstatement responding was normalized to the mean of the last three FR3 sessions, the extinction group reinstated only 44.25 ± 7.54% of acquisition responding, whereas controls reinstated 117.23 ± 21.94% (Fig. 1D). These findings demonstrate that a history of operant extinction training suppresses cue-induced reinstatement of alcohol seeking.

### Extinction reduced reactivation of acquisition-recruited dMSN ensembles during reinstatement

Having established that extinction suppresses relapse behavior, we next sought to identify the underlying cellular mechanism. Our recent work showed that dorsomedial striatal direct-pathway medium spiny neurons (dMSNs) recruited during acquisition of alcohol self-administration form ensembles that encode alcohol-related memories and their reactivation drives relapse^24^. We therefore hypothesized that extinction training suppresses reinstatement by reducing reactivation of these acquisition-recruited dMSN ensembles.

To test this, we generated triple transgenic mice (FosTRAP; Snap25-GFP; D1-tdTomato) to label neurons activated during acquisition with GFP and identify dMSNs with tdTomato (Fig. 2A). Neurons co-expressing GFP and tdTomato were defined as acquisition-tagged dMSNs (dMSN^tag^). Mice were trained for ∼8 weeks, and tagging was induced during the final two FR3 sessions via 4-hydroxytamoxifen (4-OHT; Fig. 2B, 2C) injection. Mice then underwent either 9 days of extinction or no-extinction control procedures, followed by cue-induced reinstatement testing. Ninety minutes after reinstatement, brains were collected for c-Fos immunohistochemistry to assess reactivation of acquisition-tagged dMSNs (GFP⁺ and tdTomato⁺ and c-Fos⁺).

**Figure 2.**
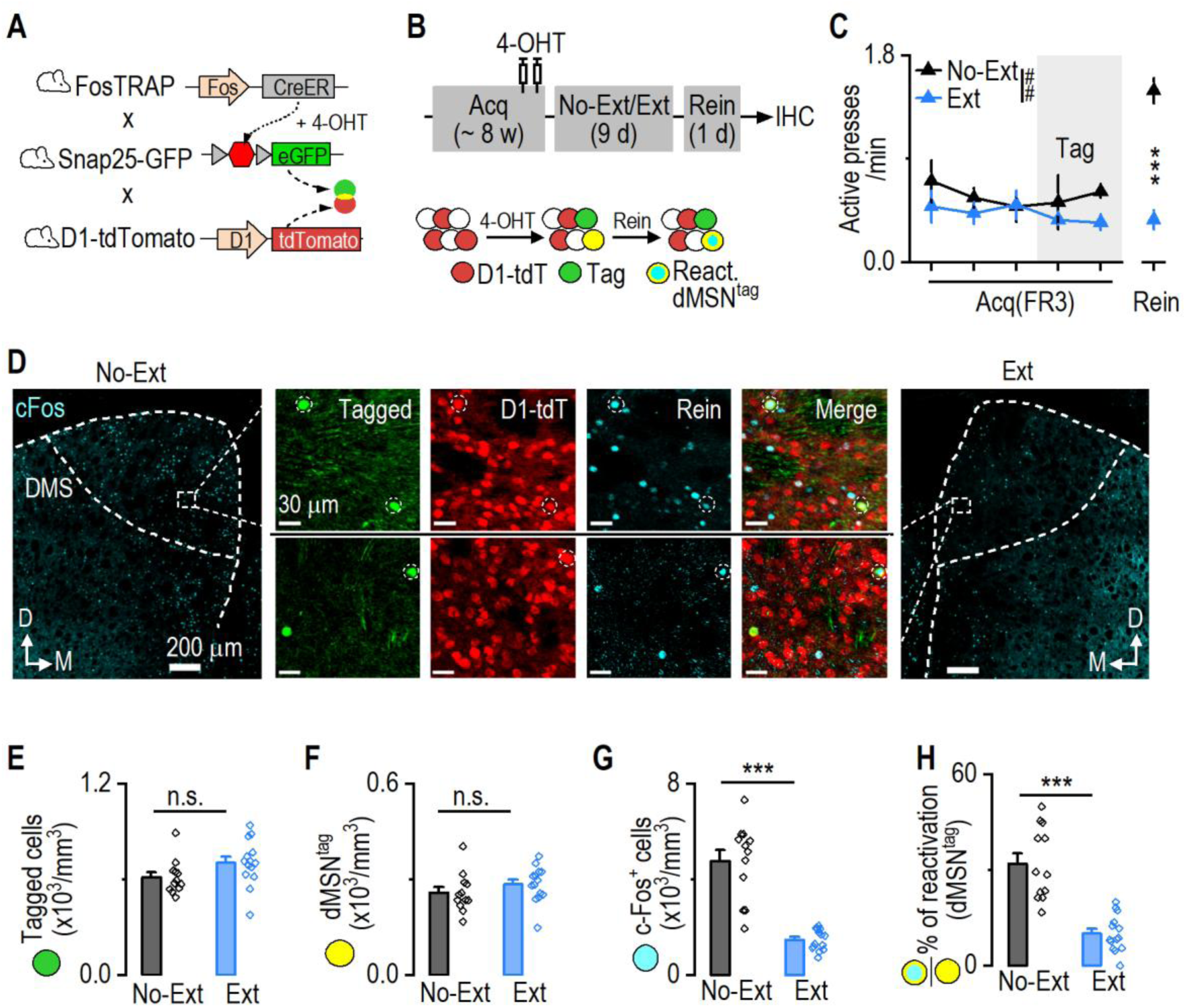
Extinction reduced reactivation of acquisition-recruited dMSN ensembles during reinstatement. ***A*,** Mechanism of activity-dependent GFP expression in tagged neurons and tdTomato (tdT) expression in dMSNs of FosTRAP;Snap25-GFP;D1-tdT mice. ***B*,** Behavioral and tagging timeline illustrating acquisition, extinction or no-extinction, reinstatement testing, and tissue collection for ensemble reactivation analysis using c-Fos immunohistochemistry (IHC). dMSNs express tdT. After tagging, tagged dMSNs (dMSN^tag^) co-express Snap25-GFP and D1-tdT. During reinstatement, reactivated dMSN^tag^ were labeled with c-Fos IHC. ***C*,** Active lever pressing during final FR3 session and reinstatement in extinction and no-extinction groups. Two-way RM ANOVA: F_(5, 20)_ = 4.981, ^##^p=0.0040; Sidak’s multiple comparisons test for No-Ext vs Ext: Acq sessions 1-5: t = 1.24, 0.74, 0.083, 0.85, 1.51, p = 0.78, 0.98, 0.99, 0.95, 0.61; Rein: t = 6.318, ***p < 0.001. ***D*,** Representative confocal images of dorsomedial striatum showing tagged acquisition neurons (GFP), D1-tdT labeling, and c-Fos expression following reinstatement. Insets show higher magnification views. ***E*,** Both groups had similar densities of tagged neurons in the dorsomedial striatum. Unpaired t test: t_(24)_ = 1.80, p = 0.08. ***F*,** Both groups had similar densities of dMSN^tag^. Unpaired t test: t_(24)_ = 1.18, p = 0.25. ***G*,** Extinction training reduced the density of reactivated dMSN^tag^ during reinstatement. Unpaired t test: t_(24)_ = 7.43, ***p < 0.001. ***H***, Extinction training reduced the percentage of acquisition-recruited dMSNs reactivated during reinstatement. Unpaired t test: t_(24)_ = 6.33, ***p < 0.001. n = 3 mice per group (C), 12 slices/3 mice (No-Ext; E-H) and 14 slices/3 mice (Ext; E-H).

Both groups displayed comparable acquisition performance (Fig. 2C). As expected, reinstatement responding was reduced in the extinction group. The density of tagged neurons and tagged dMSNs did not differ between groups (Fig. 2D-2F), confirming equivalent initial labeling during acquisition. However, extinction markedly reduced the overall number of c-Fos⁺ neurons during reinstatement (Fig. 2G), and significantly decreased the reactivation rate of tagged dMSNs [(c-Fos⁺& GFP⁺ & tdTomato⁺)/(GFP⁺ & tdTomato⁺)] (Fig. 2H).

Thus, extinction training suppresses relapse behavior in parallel with reduced reactivation of alcohol-memory-encoding dMSN ensembles, suggesting that extinction dampens the re-engagement of acquisition ensembles during relapse.

### Chemogenetic activation of acquisition-recruited dMSN ensembles during extinction impaired extinction and promoted reinstatement

If extinction suppresses relapse by reducing acquisition-ensemble activity, then artificially activating these ensembles during extinction should interfere with extinction learning and promote relapse (Fig. 3A).

**Figure 3.**
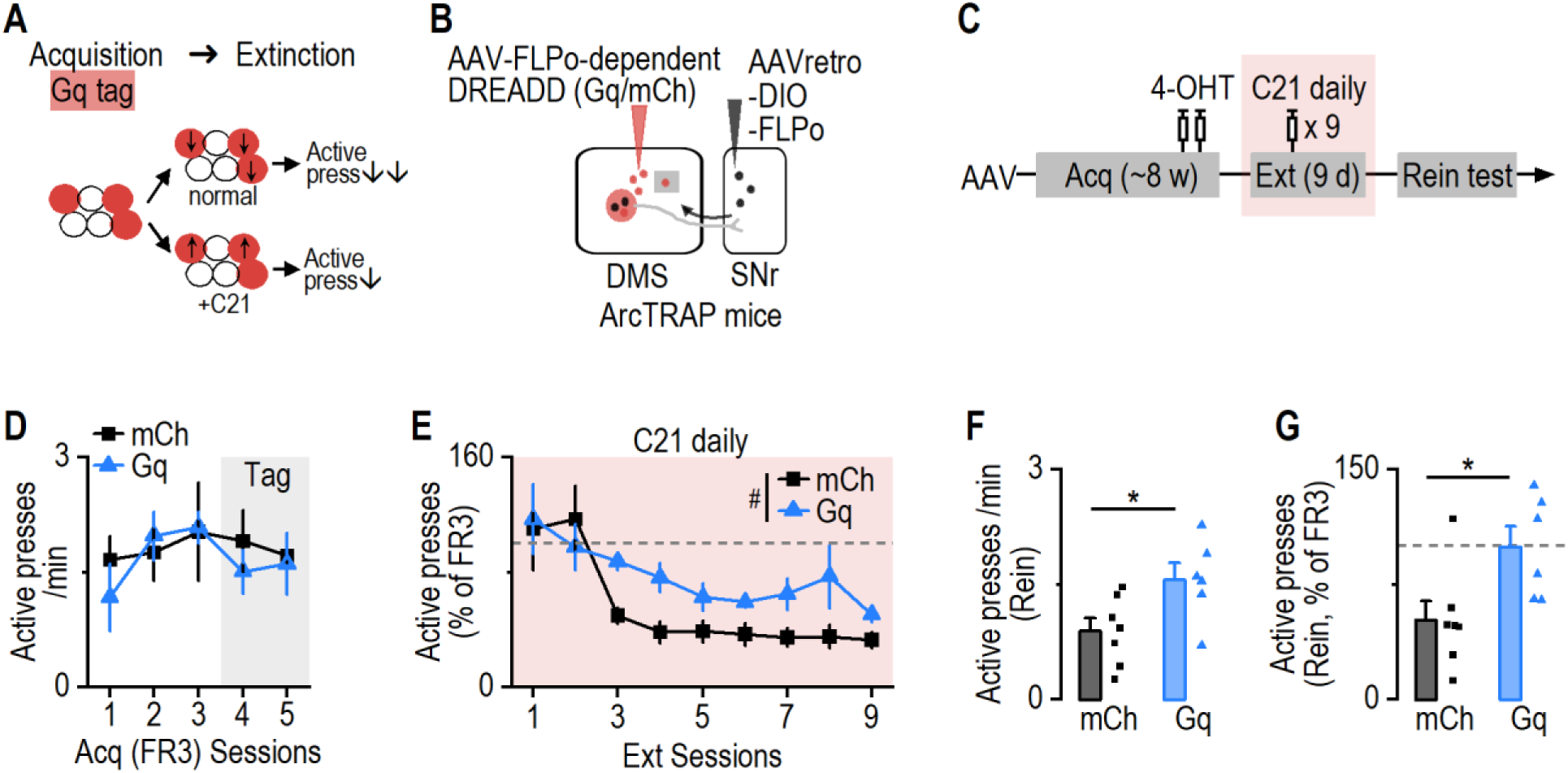
Chemogenetic activation of acquisition-recruited dMSN ensembles during extinction impaired extinction and promoted reinstatement. ***A*,** During extinction, acquisition-recruited striatal ensembles are suppressed, contributing to the reduction of alcohol-seeking behavior. We hypothesized that chemogenetic activation of these acquisition-recruited neurons during extinction would counteract this suppression, impair extinction learning, and promote relapse. ***B*,** Schematic illustrating intersectional viral approach to express DREADD in acquisition-recruited striatal dMSNs in ArcTRAP mice. ***C*,** Following viral infusion, mice underwent ∼8 weeks of operant alcohol self-administration, with tagging occurred during the last two days of the training. Two weeks later, they underwent daily chemogenetic activation (via C21 injection, 1 mg/kg) during extinction sessions, followed by cue-induced reinstatement testing without C21 injection. ***D*,** Active lever pressing during acquisition was comparable between groups. Two-way RM ANOVA: F_(1, 11)_ = 0.29, p = 0.60. ***E***, Chemogenetic activation of acquisition-recruited dMSNs resulted in persistently elevated responding across extinction sessions. Two-way RM ANOVA: F_(1, 11)_ = 5.697, ^#^p = 0.036. ***F*,** Activation of acquisition-recruited dMSNs during extinction enhanced active presses during cue-induced reinstatement. Unpaired t test: t_(11)_ = 2.46, *p = 0.032. ***G*,** Reinstatement responding expressed as a percentage of acquisition responding demonstrates that activation of acquisition ensembles during extinction promotes relapse. Unpaired t test: t_(11)_ = 2.59, *p = 0.025. n = 7 mice (mCh) and 6 mice (Gq).

To selectively manipulate acquisition-recruited dMSNs, we used ArcTRAP mice combined with a projection- and recombinase-based viral strategy (Fig. 3B)^24^. AAVretro-DIO-FLPo was injected into the substantia nigra pars reticulata (SNr), enabling retrograde labeling of dMSNs with DIO-FLPo. FLPo-dependent DREADDs were infused into the dorsomedial striatum (DMS). In combination with ArcTRAP-mediated CreER activation during acquisition via 4-OHT administration, which allowed acquisition-recruited dMSNs expressing FLPo, this strategy restricted hM3Dq (Gq) or mCherry (mCh) expression to acquisition-recruited dMSNs.

After stable FR3 performance and tagging during the final acquisition sessions, mice were allowed three weeks for viral expression before extinction training. During the 9-day extinction phase, mice received daily C21 injections (1 mg/kg, i.p.) prior to sessions to activate Gq-expressing neurons (Fig. 3B, 3C). Chemogenetic activation of acquisition-recruited dMSNs significantly impaired extinction learning, as Gq-expressing mice exhibited higher active lever pressing during the middle-to-late extinction sessions compared to controls (Fig. 3E). During a subsequent reinstatement test conducted without C21, Gq mice showed markedly elevated active pressing (Fig. 3F). When normalized to acquisition responding, Gq mice reinstated nearly 100% of acquisition levels (99.49 ± 13.49%), whereas control mice reinstated approximately half (51.91 ± 12.45%; Fig. 3G).

Thus, artificial activation of acquisition-recruited dMSNs during extinction interferes with extinction learning and consequently enhances later reinstatement, supporting with the idea that extinction normally engages mechanisms that suppress acquisition-ensemble activity.

### Chemogenetic activation of striosomal dMSNs during extinction training facilitated extinction and suppressed reinstatement

In addition to requiring suppression of acquisition-recruited dMSN ensembles to weaken alcohol-associated memories, extinction is itself considered a form of new learning. Emerging evidence suggests that extinction may preferentially recruit striosome-enriched dMSNs within the striatum^24^. We therefore hypothesized that enhancing striosomal dMSN activity during extinction would strengthen extinction learning and further reduce subsequent relapse (Fig. 4A).

**Figure 4.**
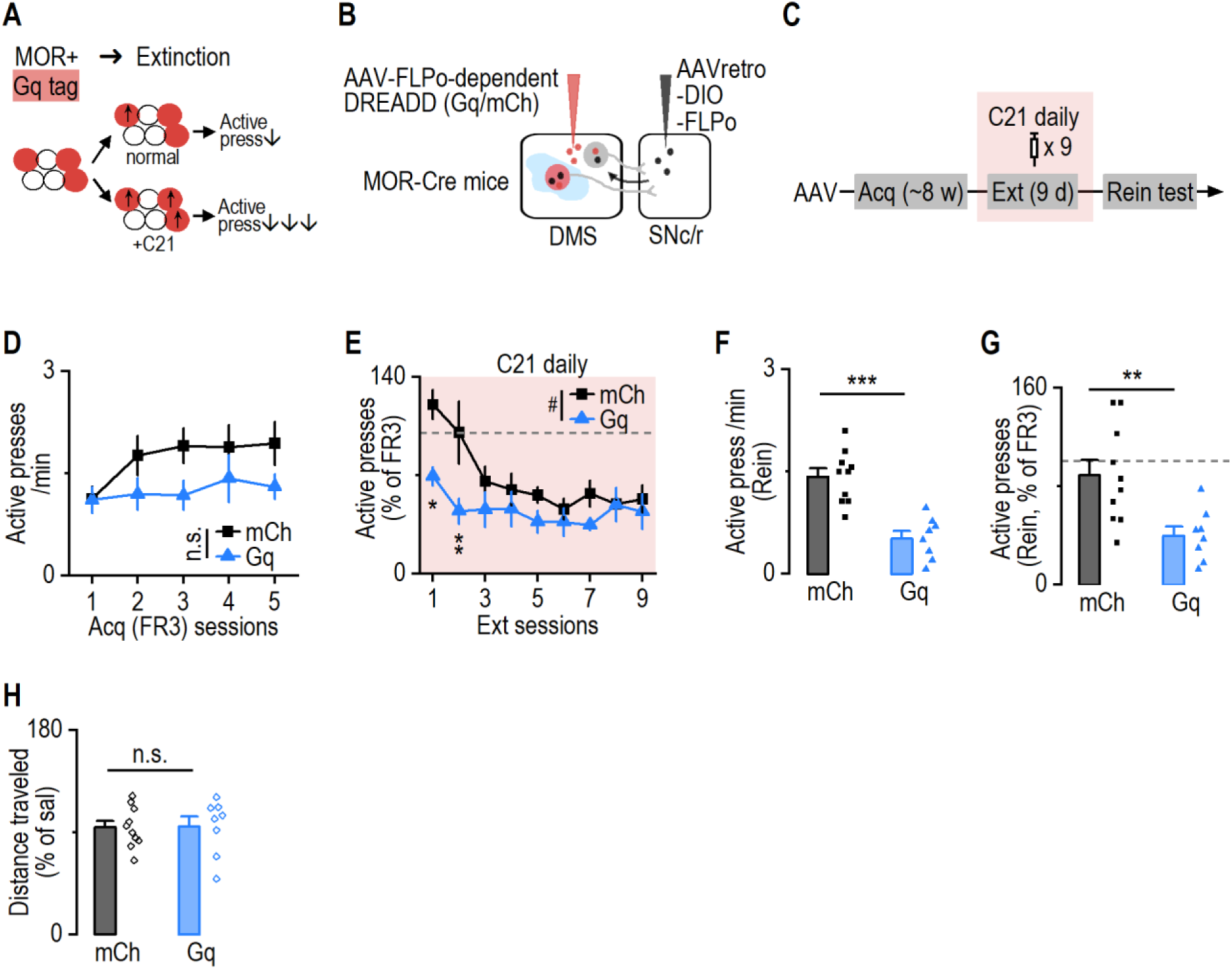
Chemogenetic activation of striosomal dMSNs during extinction training facilitated extinction and suppressed reinstatement. ***A*,** Under normal extinction conditions, striosomal dMSNs are recruited to suppress alcohol-seeking behavior, resulting in reduced active lever pressing. We hypothesized that chemogenetic activation of striosomal dMSNs during extinction would further enhance this suppression, thereby accelerating extinction learning and reducing subsequent reinstatement. ***B*,** Schematic illustrating intersectional viral approach to express DREADD in striosomal dMSNs in MOR-Cre mice. ***C*,** Following acquisition and viral expression, mice underwent daily chemogenetic activation during extinction sessions, followed by reinstatement testing. ***D,*** Active lever pressing during acquisition was comparable between groups. Two-way RM ANOVA: F_(1, 16)_ = 2.28, p = 0.15. ***E*,** Activation of striosomal dMSNs accelerated extinction learning and reduced responding across extinction sessions. Two-way RM ANOVA: Group × Session: F_(8, 128)_ = 2.41, ^#^p = 0.0185. Sidak’s multiple comparisons test for mCh vs Gq: session 1: t = 3.23, *p = 0.014; session 2: t = 3.53, **p = 0.0049. ***F*,** Activation of striosomal dMSNs during extinction reduced active presses during reinstatement. Unpaired t test: t_(16)_ = 5.39, ***p < 0.001. ***G,*** Reinstatement responding expressed as a percentage of acquisition responding demonstrates that activation of striosomal dMSNs during extinction suppressed relapse. Unpaired t test: t_(16)_ = 3.5, **p = 0.0029. ***H***, Chemogenetic activation of striosomal dMSNs did not alter distance traveled in an open field test. Unpaired t test: t_(16)_ = 0.065, p = 0.95. n = 10 mice (mCh) and 8 mice (Gq).

To selectively target striosomal dMSNs, we used MOR-Cre mice in combination with the same intersectional dual-virus strategy (Fig. 4B). AAVretro-DIO-FLPo was infused into the SNc/SNr, enabling retrograde labeling of striatal dMSNs. Because FLPo expression was Cre-dependent, only MOR-expressing (striosomal) dMSNs expressed FLPo, while other dMSNs were excluded. Subsequent infusion of an FLPo-dependent hM3Dq (Gq) or mCherry virus into the DMS restricted DREADD expression specifically to striosomal dMSNs. After mice achieved stable performance under the FR3 schedule (Fig. 4C, 4D), they underwent 9 days of extinction training with daily C21 administration to activate Gq-expressing striosomal dMSNs.

Chemogenetic activation of striosomal dMSNs significantly accelerated extinction, particularly during the early sessions, as Gq mice displayed reduced active lever pressing compared to controls (Fig. 4E). During the subsequent cue-induced reinstatement test (without C21), Gq-expressing mice showed significantly lower active responding and reduced normalized reinstatement compared to controls (Fig. 4F, 4G). Importantly, activation of striosomal dMSNs did not alter locomotor activity in an open-field test (Fig. 4H), indicating that the behavioral effects were not attributable to nonspecific motor changes.

Together, these findings demonstrate that enhancing striosomal dMSN activity during extinction facilitates extinction learning and strengthens extinction memory, resulting in more persistent suppression of relapse. These results suggest that extinction training suppresses relapse through coordinated mechanisms that both reduce the activity of acquisition-recruited dMSN ensembles and actively engage striosomal dMSNs to support extinction learning.

## Discussion

In this study, we identify coordinated striatal mechanisms through which extinction suppresses cue-induced reinstatement of alcohol seeking. We show that extinction reduces reactivation of acquisition-recruited dorsomedial striatal dMSN ensembles during reinstatement, that preventing suppression of these ensembles during extinction impairs extinction learning and enhances relapse, and that enhancing striosomal dMSN activity during extinction facilitates extinction learning and produces more persistent suppression of relapse. Together, these findings indicate that extinction limits alcohol seeking by rebalancing striatal dynamics—dampening previously established alcohol-memory representations while engaging specific striatal subpopulations to support updated behavioral control.

### Extinction suppresses reactivation of acquisition-recruited dMSN ensembles

A central finding of this study is that extinction training reduces reactivation of acquisition-recruited dMSN ensembles during cue-induced reinstatement. Prior work demonstrated that dMSNs recruited during operant alcohol self-administration encode alcohol-associated memories and that their reactivation drives relapse-like behavior^24^. However, whether extinction modifies the subsequent engagement of these ensembles remains unclear. Our data show that extinction significantly decreases reactivation of acquisition-tagged dMSNs during reinstatement, in parallel with reduced alcohol-seeking behavior. These findings suggest that extinction attenuates the behavioral influence of previously established alcohol-memory ensembles.

Importantly, chemogenetic activation of acquisition-recruited dMSNs during extinction impaired extinction learning and resulted in elevated reinstatement, even though DREADD activation was absent during the reinstatement test. This dissociation indicates that maintaining acquisition-ensemble activity during extinction does not simply enhance alcohol seeking acutely, but interferes with the formation or consolidation of extinction memory. Notably, activation of acquisition ensembles did not increase active lever pressing at the beginning of extinction, when extinction memory had not yet formed. This suggests that early in extinction, suppression of acquisition-ensemble activity has not yet become a dominant regulatory process. As extinction progresses, however, endogenous suppression of acquisition ensembles appears to emerge, and artificial reactivation of these neurons disrupts this process. The delayed extinction observed in Gq-expressing mice therefore likely reflects interference with extinction-dependent suppression mechanisms rather than a direct enhancement of alcohol-seeking behavior. Together, these findings indicate that suppression of acquisition-recruited dMSN activity is necessary for effective extinction learning.

These results also inform the longstanding debate regarding whether extinction erases prior memories or instead establishes an inhibitory trace. Our data do not support memory erasure. Acquisition-tagged neurons remain anatomically intact and capable of reactivation. Rather, extinction appears to reduce the probability that these ensembles are re-engaged during relapse. Numerous studies indicate that memory is encoded through ensemble-specific synaptic potentiation^41,42^, most prominently at glutamatergic synapses onto defined neuronal populations. Extinction may therefore operate, at least in part, by partially reversing or functionally weakening such synaptic potentiation^43–45^, thereby reducing the efficacy of presynaptic inputs in driving postsynaptic ensemble activation. Ensemble excitability state is a key determinant of memory retrievability^46^. By modulating synaptic strength and/or intrinsic excitability within acquisition-recruited dMSNs, extinction could decrease the likelihood that alcohol-associated cues successfully recruit these ensembles. In this framework, extinction reshapes the functional accessibility or dominance of alcohol-memory ensembles within striatal circuitry, limiting their behavioral expression without eliminating the underlying memory representation.

### Extinction as active striatal learning: recruitment of striosomal dMSNs

In addition to suppressing acquisition-encoded ensembles, extinction is widely viewed as an active learning process that encodes updated action–outcome contingencies^2,11^. Emerging evidence suggests that striosomal neurons are preferentially engaged during behavioral updating and decision conflict^29,31,47,48^, raising the possibility that they contribute to extinction learning. Our findings support this idea. Chemogenetic activation of striosomal dMSNs during extinction accelerated the reduction of active lever pressing and resulted in significantly lower reinstatement during subsequent cue exposure. Notably, these effects persisted in the absence of DREADD activation during reinstatement, indicating that enhanced striosomal activity strengthened extinction memory rather than transiently suppressing behavior. Moreover, the lack of changes in locomotor activity suggests that these effects reflect specific modulation of alcohol-seeking circuits rather than nonspecific motor suppression.

Striosomal dMSNs are uniquely positioned to influence learning and behavioral updating. They receive preferential limbic cortical input and project directly to dopaminergic neurons in the SNc^30,49,50^, forming a circuit that can modulate dopaminergic signaling. Through this architecture, striosomal activity may regulate prediction error signals and value updating during extinction. During extinction learning, limbic cortical inputs could recruit striosomal dMSNs, which in turn modulate SNc dopamine neuron activity. Our recent work showed that extinction training reduced striatal dopamine transients during reinstatement^5^, which might be due to striosomal dMSN activation. By shaping dopaminergic signaling, striosomes may influence extinction-related teaching signals, thereby facilitating the encoding of updated contingencies in which lever-press no longer predict reward. Through this mechanism, striosomal engagement may actively support the formation and stabilization of extinction memory. Beyond local striatal effects, modulation of dopaminergic output by striosomal dMSNs could also influence motivational circuits downstream of the midbrain, further contributing to reduced relapse vulnerability.

### Coordinated striatal mechanisms underlying extinction-dependent suppression of relapse

Taken together, our findings suggest that extinction suppresses relapse through coordinated modulation of distinct striatal dMSN populations. Successful extinction appears to require both reduction of acquisition-ensemble influence and engagement of striosomal circuits that promote behavioral updating. Rather than representing independent or competing memory traces in isolation, these processes likely interact dynamically within striatal networks to rebalance action selection in favor of abstinence.

This framework refines prevailing models of relapse and extinction. Relapse vulnerability may arise when acquisition-encoded ensembles retain dominant influence over striatal output, allowing alcohol-associated cues to drive behavior. Extinction training may reduce relapse risk by shifting ensemble dynamics—decreasing the likelihood that acquisition-recruited dMSNs are reactivated while strengthening striosomal contributions that encode updated contingencies. Importantly, these findings highlight that extinction is not only mediated by cortical, hippocampal, or amygdalar circuits but also involves intrinsic reorganization within the dorsal striatum. Our results demonstrate that extinction reshapes the activity of defined striatal projection neuron populations within this region, providing a cellular substrate for the suppression of alcohol seeking.

### Implications for addiction and therapeutic strategies

Persistent relapse remains a hallmark of alcohol use disorder, and extinction-based behavioral therapies exhibit variable efficacy. The present findings suggest that variability in extinction success may reflect differential engagement of striatal ensemble dynamics. Insufficient suppression of acquisition-recruited ensembles or inadequate recruitment of striosomal circuits could leave individuals vulnerable to cue-induced relapse. Interventions that selectively enhance extinction-related striatal plasticity or reduce the influence of maladaptive acquisition ensembles may therefore improve therapeutic outcomes.

In summary, extinction training suppresses alcohol relapse through coordinated striatal mechanisms: attenuating reactivation of acquisition-recruited dMSN ensembles and engaging striosomal dMSNs to support extinction learning. These findings establish a circuit-level framework for understanding how extinction reshapes striatal ensemble dynamics to limit relapse and provide a foundation for developing targeted strategies to enhance recovery from addiction.

## Materials and Methods

### Animals

Drd1a-tdTomato (#016204), Snap25-LSL-F2A-GFP (#021879), FosTRAP (Fos^2A-iCreER^; #030323), and ArcTRAP (Arc^CreER^; #021881)^37,51,52^ mice were purchased from the Jackson Laboratory. MOR-Cre were given by Dr. Brigitte Kieffer. FosTRAP mice were crossed with Snap25-GFP and D1-tdTomato mice to obtain FosTRAP;Snap25-GFP;D1-tdTomato mice. Genotypes were determined by PCR analysis of Cre or the fluorescent protein gene in tail DNA. All animals that were used are in mixed gender and aged from 3-8 months, and they were randomly assigned to experimental groups. Animals were housed in a temperature- and humidity-controlled vivarium with a reversed 12-h light/dark cycle (lights on at 10:00 P.M. and off at 10:00 A.M.). All behavior experiments were conducted in their dark cycle, starting approximately 1 h after the light went off. Food and water were available *ad libitum*. All animal care and experimental procedures were approved by the Texas A&M University Institutional Animal Care and Use Committee.

## Method Details

### Intermittent access to 20% alcohol two-bottle-choice drinking procedure

To establish high levels of alcohol consumption in mice, we utilized the intermittent-access to 20% alcohol two-bottle-choice drinking procedure^16,19,22,53–57^. Mice were given free access to two bottles containing water or 20% alcohol for three 24-h sessions (Mondays, Wednesdays, and Fridays), with 24-h or 48-h withdrawal periods (Tuesdays, Thursdays, Saturdays, and Sundays) each week. During the withdrawal periods, the mice had unlimited access to water bottles. The placement of the alcohol bottle was alternated for each drinking session to control side preferences. For all animals that underwent operant self-administration of alcohol, 3-6 weeks of 2BC were used to acclimate them with alcohol.

### Operant self-administration of alcohol

Animals were trained to self-administer 20% alcohol in operant chambers using a fixed ratio (FR) schedule^58–60^. The mouse operant setup (Med Associates) included two levers in each chamber: active and inactive, on the opposite sides of the same wall.

During alcohol OSA training sessions, a house light centered above the operant chamber was illuminated. Each operant experiment commenced with continuous reinforcement (FR1) every other day for approximately two to three weeks, wherein pressing the active lever resulted in a ∼0.32 mL delivery of 20% alcohol. Actions on the inactive side were documented but did not initiate a programmed event. The alcohol solution was dispensed into a stainless-steel reservoir situated within the magazine port between active and inactive levers. Each alcohol delivery persisted for 3 sec and was accompanied by a tone and a discrete yellow cue light in the magazine port for the same duration. There was a 20-sec time-off period for each delivery, during which active/inactive presses were recorded but did not cause any further alcohol deliveries. When animals were able to achieve at least 10 deliveries under the FR1 schedule, the criterion to receive the alcohol escalated to FR2 (around one to two weeks), and eventually progressed to FR3 (around two weeks). Each operant session for mice lasted 30 min or 1 h, depending on the learning condition.

#### Extinction training

The extinction training was conducted under the FR3 schedule. However, pressing on the active side did not result in actual alcohol delivery, nor cue presentation. Each extinction session for mice lasted 30 min, and in total lasted for 9 d. For chemogenetic manipulations during the extinction, C21 (i.p., 1 mg/kg; or saline) was injected 30 min prior to the start of the test session.

#### Cue-induced reinstatement of alcohol seeking

The reinstatement test (30 min) was conducted as previously described^61^. It was carried out at the FR3 schedule, where three actions on the active side triggered the illumination of the cue light within the magazine port, the tone, and the pump action sound, with the exception that the first cue presentation was given immediately after the first active press.

### Tag active neurons

4-OHT was prepared and tagging was conducted as previously described^37,52,62^. The 4-OHT solution was freshly prepared on the tagging day. In brief, 4-OHT powder was dissolved in 200 proof ethanol (20 mg/mL) at room temperature, with continuous shaking until the 4-OHT was fully dissolved. An equal volume of a 1:4 mixture of castor oil and sunflower seed oil was added, resulting in a final concentration of 10 mg/mL 4-OHT.The ethanol was removed by centrifugation under vacuum. The final 4-OHT in oil was stored at 4°C for a maximum of 12 h before use.

For tagging during acquisition of operant learning, mice were habituated to i.p. injection of saline immediately after each operant session for two consecutive days. On the third and fourth operant sessions (Tag), mice were introduced to an isoflurane chamber immediately after the operant session and 4-OHT was i.p. injected. The use of isoflurane is to reduce the pain and discomfort due to 4-OHT/oil mix injection. After the injection, mice were returned to their home-cage and left without any disturbance for the following 6 hours.

### Stereotaxic virus infusion and fiber implantation

Where required for the experimental design, AAVretro-DIO-FLPo (0.6 µL/site) was bilaterally infused into the SNr (AP: -3.15 mm; ML: ±1.35 mm; DV: -4.6 mm), and AAV-fDIO-mCherry/AAV-fDIO-hM3Dq-mCherry (0.5 µL/site) was bilaterally infused into the DMS (AP1: +0.98 mm; ML1: ±1.25 mm; DV1: -2.9 to -3.1 mm; AP2: +0.3 mm; ML2: ±1.87 mm; DV2: -2.9 to -3.1 mm) of ArcTRAP or MOR-Cre mice. Animals were anesthetized with 3-4% isoflurane at 1.0 L/min and mounted in a stereotaxic surgery frame. The head was leveled and craniotomy was performed using stereotaxic coordinates adapted from the brain atlas ^63^. The virus was infused at a rate of 0.08 µL/min. At the end of the infusion, the injectors remained at the injection site for an additional 10-15 min before removal to allow for virus diffusion. The scalp incision was then sutured, and the animals were returned to their home cage for recovery.

### Open field locomotor activity test

A transparent open field activity chamber (Med Associates, 43 cm x 43 cm x 21 cm height) was equipped with an infrared beam detector that was connected to a computer. All tests lasted for 10 min. Animals were first habituated in the chamber to remove the novelty-induced effect. Then, animals were tested for locomotor activity by placing them into the chamber 30 min after i.p. injections of saline. One day later, animals were re-introduced to the chamber 30 min after i.p. injections of C21 (1 mg/kg). Locomotor activity during the C21-injected day was compared between groups by normalizing the distance traveled in the C21-injected day to the distance travelled in the saline-injected day.

### Histology and cell counting

Animals were perfused intracardially with 4% formaldehyde in phosphate-buffered saline (PBS). The brains were removed and post-fixed overnight in 4% formaldehyde in PBS at 4°C prior to dehydration in 30% sucrose solution. The brains were then cut into 50-µm coronal sections using a cryostat. c-Fos IHC was carried out on free-floating sections. All incubations (except for the primary antibody) and rinses took place at room temperature on a shaker. Sections were rinsed in PBS three times between steps. Sections were first placed in blocking solution with 10% bovine serum albumin (BSA) in PBS with 0.3% triton-X (PBST) for 1 hour and incubation in primary antibody (rabbit anti-c-Fos: for mice: 1:2000, EMD-Millipore, #PC38) diluted in blocking solution (1% BSA in 0.3% PBST) overnight under 4°C. Sections were then incubated in biotinylated donkey anti-rabbit (1:500, Jackson Immuno Research) in blocking solution for 1 hour. Sections were visualized via incubation for 1 hour in Streptavidin-conjugated Alexa 647 (1:1000, Thermo Fisher) in PBS. After incubation, all slices were washed 3 times in PBS and mounted.

All images were acquired using a confocal microscope (FluoView 3000, Olympus, Tokyo, Japan) and analyzed using IMARIS 8.3.1 (Bitplane, Zürich, Switzerland), as previously reported^18,64^. For cell counting in slices from FosTRAP mice, analyses were performed within a circular region-of-interest (ROI; radius = 300 µm) positioned consistently at the dorsal-medial boundary of the DMS across all slices.

### Quantification and Statistical analysis

All data are presented as mean ± standard error of the mean (SEM). Normality was assessed using the Shapiro-Wilk test. If the data met normality assumptions, statistical comparisons were conducted using a two-tailed t-test (paired or unpaired) or one-, or two-ANOVA (with repeated measures), followed by post hoc Tukey’s or Sidak’s multiple comparisons test, when appropriate. If normality was not met, data were analyzed using the Mann-Whitney test, Kruskal-Wallis test (followed by Dunn’s post hoc test), or mixed-effects analysis with post hoc Tukey’s or Sidak’s multiple comparisons test. Statistical significance was set at p < 0.05. All statistical analyses were performed using SigmaPlot, and graphs were generated using OriginPro.

## Acknowledgements

We thank the National Institute on Mental Health Drug supply program for giving 4-OHT and C21. This research was supported by NIAAA Grant R01AA021505, R01AA027768, U01AA025932, R01AA030293 and X-grant from Texas A&M University to J.W., McGovern Fellowship from Texas Research Society on Alcoholism (TRSA) to X.X., and Doctoral Student Small Grant from Research Society on Alcohol (RSA) to X.X.

## Author contributions

Conceptualization: J.W. and X.X.; Methodology: X.X. and J.W.; Investigation and Formal analysis: X.X.; Writing - Original Draft: X.X.; Writing – Review & Editing: J.W. and X.X.; Resources: X.W.; Funding acquisition: J.W. and X.X.; Supervision: J.W.

## Declaration of interests

The authors declare no competing interests.

## Resource availability

### Lead contact

Requests for further information and resources should be directed to and will be fulfilled by the lead contact, Jun Wang.

### Materials availability

This study did not generate new unique reagents.

### Data and code availability

- All data reported in this paper will be shared by the lead contact upon request.
- This paper does not report original code.
- Any additional information required to reanalyze the data reported in this paper is available from the lead contact upon request.

### Declaration of generative AI and AI-assisted technologies in the writing process

During the preparation of this work, the authors used ChatGPT (OpenAI) to improve the grammar and readability of the manuscript. After using this tool, the authors reviewed and edited the content as needed and took full responsibility for the content of the published article.

## Notes

### Competing Interest Statement

The authors have declared no competing interest.

